# Do shifts in honeybee crop microbiota enable ethanol accumulation? A comparative analysis of caged and foraging bees

**DOI:** 10.1101/2025.06.05.658140

**Authors:** Weronika Antoł, Bartłomiej Surmacz, Monika Ostap-Chec, Daniel Stec, Krzysztof Miler

**Affiliations:** Institute of Systematics and Evolution of Animals of the Polish Academy of Sciences, Kraków, Poland

**Keywords:** foregut, caged bees, microbiome, fermentation, auto-brewery, alcohol

## Abstract

Honeybees encounter low environmental doses of ethanol, primarily through fermenting nectar, which can have both beneficial and detrimental effects on their functioning. Yet, ethanol traces can also be detected in the crop of caged bees with no access to environmental food sources. This raises the possibility that endogenous ethanol accumulation could occur under restricted conditions, with microbial contributions as a potential mechanism. The crop microbiota, although less diverse than that in other gut segments, plays important roles in food fermentation and pathogen defense. We hypothesized that captivity-induced shifts in crop microbiota may facilitate fermentation, resulting in measurable ethanol. To test this, we compared the crop contents of naturally foraging hive bees and caged bees reared without access to the natural environment. Ethanol levels were low in both groups and did not differ significantly, but non-zero measurements were more frequently observed in caged bees. Microbial community structure differed strongly in α- and β-diversity. Caged bees showed reduced abundance of nectar-associated genera (e.g., *Apilactobacillus*) and an increase in genera that include known ethanol-producing strains, such as *Gilliamella* and *Bifidobacterium*. While we did not directly assess metabolic activity, our results suggest that captivity alters microbial communities in ways that may influence ethanol levels. This raises broader questions about how microbe-host interactions modulate host phenotypes under different environmental conditions.

## Introduction

Ethanol is a naturally occurring substance in fermenting nectar and fruit, making it a frequent diet component for many nectar- and fruit-consuming animals [1]. In insects, ethanol often functions as an attractant and may be treated as a cue for sugar-rich food or an energy source [2]. It has been shown to attract beetles [3, 4], butterflies [5], fruit flies [6], and bees [7, 8]. In some insects, such as the Oriental hornet *(Vespa orientalis)*, ethanol is metabolized with extraordinary efficiency, even at high concentrations [9]. In *Drosophila melanogaster*, ethanol-rich environments are preferred for egg-laying [6], and fruit flies can use ethanol as a self-medication agent [10].

Honeybees are no exception to this pattern. Ethanol serves as a substrate for the synthesis of ethyl oleate [11], a pheromone regulating worker maturation into foragers [12]. Laboratory studies have shown that honeybees willingly consume sugar solutions containing up to 2.5% ethanol, sometimes even preferring them to pure sucrose [8]. Honeybee workers possess alcohol dehydrogenase (ADH), an enzyme contributing to ethanol metabolism [13], and recent findings suggest that their ethanol intake increases in response to parasitic infection, indicating a potential role in infection-induced changes in behavior [14]. Even though prolonged ethanol consumption appears to exert toxic effects in honeybees [8, 15], this effect is observed only in older bees, with no detectable impact on the survival or physiological condition of relatively young workers [16]. Honeybees and possibly other insect pollinators are likely adapted to ethanol consumption [17].

One key site of ethanol processing in honeybees is the crop, a section of the foregut that functions as a nectar reservoir and site of ethyl oleate production [18]. The crop itself does not absorb the nutrients, but its contents are selectively passed to the midgut via the proventriculus [19, 20]. Passage rate depends on the substance and the physiological state of the bee. For example, glucose is transported more slowly in immobilized than in moving bees, and the emptying rate decreases with increasing solution molarity [19]. Incidentally, we detected traces of ethanol in the crop contents of caged bees that had no access to environmental food sources. This raises the possibility that microbial activity could contribute to crop ethanol accumulation in the absence of environmental ethanol exposure. In humans, a similar phenomenon is known as gut fermentation or auto-brewery syndrome [21], with such processes described in the urinary tract as bladder fermentation syndrome [22, 23]. Ethanol-producing microbes in the human gut include bacteria and fungi such as *Candida*, *Saccharomyces,* and *Klebsiella* [21, 24–27].

The honeybee gut microbiota is established within days of adult emergence [28] and comprises a small set of core genera [29–32] including *Gilliamella, Snodgrassella, Lactobacillus melliventris* clade*, Bombilactobacillus*, *Apilactobacillus, Bifidobacterium* and, less abundant, *Frischella, Bartonella, Bombella* and *Gluconobacter* [32–35]. These communities show large strain-level and functional diversity. All of them play significant roles in digestion, immune defence, and detoxification, and show seasonal variation [29, 32, 33, 36–38]. These bacterial communities can also protect against pathogens [39–41] and modulate the effects of pesticides [42], or even produce neuroactive metabolites, such as GABA or the juvenile hormone [43, 44]. Some of these honeybee gut bacteria, such as *Gilliamella, Bifidobacterium,* and *Bombilactobacillus* [33, 35, 45, 46], are capable of ethanol fermentation.

The honeybee microbiota is shaped by many factors, including social transmission between bees and environmental exposure via nectar, pollen, flower surfaces, and honey [47]. The crop, though less diverse than other gut regions, harbors a characteristic microbial community dominated by *Apilactobacillus kunkeei* and *Bombella apis* [18, 33]. Its composition is relatively similar to that which characterises corbicular pollen [48]. The crop microbiota participates in honey and bee bread production [49, 50], inhibits pathogens and environmental microbes [39–41], and is transmitted between bees via trophallaxis (mouth-to-mouth food exchange) [28].

In this study, we addressed two questions: (1) how does the absence of environmental exposure affect the crop microbiota of caged bees, and (2) does this shift in microbial composition potentially contribute to crop ethanol accumulation? To address these questions, we sampled the crop contents of age-matched bees kept in natural hive conditions or laboratory cages. We assessed both microbiota composition and ethanol levels after one and two weeks. We hypothesized that confinement would reduce crop bacterial diversity due to the loss of environmental inputs. We further hypothesized that this loss of environmentally derived bacteria could open ecological space for otherwise less abundant taxa, potentially creating niches for ethanol-producing and other opportunistic species in caged bees over time.

## Methods

### Experimental design

The experiment was conducted using queen-right colonies of *Apis mellifera carnica* Poll with naturally inseminated queens. All colonies were in good overall condition. The study was performed in three replicates, each separated by a 24-hour interval.

For each replicate, newly emerged bees were obtained from two unrelated colonies. To achieve this, a brood frame with capped cells, free of adult bees, was collected from each colony and placed overnight in an incubator (KB53, Binder, Germany) set at 32 °C. The following morning, all newly emerged bees were marked with a colored dot on the thorax using a non-toxic paint marker and introduced into an unrelated hive, allowing them to develop naturally.

At seven days of age, a subset of the marked bees was collected from the hive using forceps, transferred to wooden cages, and transported to the laboratory. For each replicate, four cages were established, each containing 50 individuals. Throughout the laboratory phase, bees were provided with a 40% sucrose solution and water *ad libitum* via gravity feeders and maintained in an incubator (KB400, Binder, Germany) at 32 °C. Sucrose and water were replenished daily, and dead individuals were removed. The remaining marked bees in the hive continued their natural development and were collected later as needed.

The first sampling was conducted when the bees reached 14 days of age. Two out of the four cages per replicate were removed from the incubator for sampling. Simultaneously, additional marked bees of the same age were collected from the hive and placed into two separate wooden cages (50 individuals per cage). To preserve their natural gut contents, these newly collected bees were not provided with sucrose solution or water after collection. This design resulted in two groups of same-age bees with distinct rearing environments (hereafter referred to as ‘bee sources’): (1) bees that had spent the previous week in the laboratory (‘caged bees’) and (2) bees that had remained in the hive (‘hive bees’). Crop content was sampled from living bees. Within each replicate, one cage per bee source was designated for ethanol content analysis, while the other was used for microbiota metabarcoding.

The second sampling followed the same procedure after another week, when the bees reached 21 days of age. At this stage, the remaining two incubator cages from each replicate were used, along with two additional cages containing newly collected, marked bees from the hive. As before, the newly collected bees were not provided with any food or water to maintain the integrity of their natural digestive tract contents.

### Ethanol content analysis

#### Sample collection

Crop content was sampled individually from both bee sources (hive vs. caged bees) at two timepoints (week 1 vs. week 2) across three replicates (244 individuals in total). To sample crop content, individual bees were carefully removed from their cages using forceps and gently pressed against a Styrofoam plate until they regurgitated a fluid droplet. The fluid was collected into an end-to-end microcapillary tube, placed in a cryotube, and immediately frozen at −20 °C until analysis. Care was taken to avoid potential leakage from other parts of the digestive system, and visibly contaminated (opaque) samples were discarded.

Additionally, food and water samples that the caged bees had access to in the 24 hours preceding sampling, as well as freshly prepared sucrose solution and water used for feeder replenishment on the sampling day, were collected (24 samples in total). Food samples were diluted 100x before analysis as an in-house procedure in our laboratory, covering various ethanol concentrations.

#### Ethanol assay

Ethanol levels were measured using an ethanol assay kit (K-ETOH, Neogen, USA), following the manufacturer’s protocol. Three µl per sample was taken for analysis, with the remaining volume filled to the required 10 µl using distilled water. Several samples were pooled to achieve the minimum 3 µl, or discarded, if failed to achieve the required volume even after pooling, resulting in a total of 127 crop samples: 85 from hive bees (48 and 37 from week 1 and week 2, respectively) and 42 from caged bees (26 and 16 from week 1 and week 2, respectively); see Online Resource 1: Table S1 for samples summary. Absorbance was measured at 340 nm using a Multiskan FC microplate reader (Thermo Scientific, USA). Each batch (plate) included a calibration curve covering the range of 0.0125 – 0.1 g/L ethanol. Ethanol levels were calculated using these calibration curves and corrected for the sample dilution factor. Given the 3.33x dilution (3 µl of sample filled to the required 10 µl reaction volume with distilled water), the practical detection threshold for the assay was 0.033 g/L, based on the manufacturer’s minimum reliable detection limit of 0.01 g/L. Results below this threshold were considered below the detection limit and recorded as 0.

#### Data analysis

Statistical analysis was performed in R [51]. Ethanol levels were analyzed using a generalized linear mixed-effects model (glmmTMB, ziGamma with log link) fitted using the ‘glmmTMB’ package [52], including bee source, timepoint, and their interaction as fixed effects, and bee source as the zero-inflation model variable. The model with replicate had a lower fit (according to the Akaike Information Criterion), therefore, this factor was not included. Model diagnostics were performed using the ‘DHARMa’ package [53].

### Microbiota metabarcoding

#### Sample collection

Crop content was sampled from both bee sources at two timepoints across three replicates (207 individuals in total). The sampling procedure matched that used for ethanol content analysis, except that the regurgitated fluid was collected directly into sterile 0.5 ml Eppendorf tubes. To increase the sample volume and DNA concentration, samples were pooled (7-14 samples per pool, depending on individual sample volume), resulting in 11 samples from the caged bees and 12 samples from the hive bees (23 samples in total, with volumes ranging from 10 to 100 µl; see Online Resource 1: Table S1).

Food and water that the caged bees had access to in the 24 hours preceding sampling, as well as freshly prepared sucrose solution and water used for replenishing feeders on the sampling day, were also sampled (24 samples in total, 50 µl per sample).

#### DNA extraction and sequencing

DNA was extracted using the Qiagen DNeasy^®^ Blood & Tissue kit, following the manufacturer’s ‘Purification of Total DNA from Animal Tissues - Spin-Column Protocol’ and eluted in 50 µl. One extraction blank was included. DNA concentration was measured using the Invitrogen™ Qubit™ dsDNA High-Sensitivity Assay Kit.

Library preparation and sequencing were outsourced to IGA Technology Services S.R.L. (Italy). Libraries for the V3-V4 region of the *16S* rRNA gene were prepared following the Illumina 16S Metagenomic Sequencing Library Preparation protocol [54]. Universal prokaryotic primers, Pro341F and Pro805R [55], were used in the initial PCR amplification step. The subsequent amplification integrated relevant flow-cell binding domains and unique indices (NexteraXT Index Kit, FC-131-1001/FC-131-1002). Libraries were sequenced on an Aviti instrument (Element Biosciences, San Diego, USA) using a 300-bp paired-end mode. Base calling, demultiplexing, and adapter masking were performed with Bases2fastq software v.1.8 (Element Biosciences).

#### Data preprocessing

Primer sequences were identified and trimmed using Cutadapt 4.6 [56]. Subsequent data processing and analyses were performed in R [51]. The primer-trimmed reads were denoised, merged, and chimeras were removed using the ‘DADA2’ package [57], following the standard DADA2 pipeline tutorial [58] (see Online Resource 1: Table S2 for the number of reads).

#### Classification and decontamination

Taxonomy was assigned to the amplicon sequence variants (ASVs) using the IdTaxa function [59] from the ‘DECIPHER’ package [60], with the BEExact reference database (BEEx_v2023.01.30 idtaxa_v3v4.RData), which is designed for *16S* rRNA-based studies of bee-associated microbiota [61]. ASVs not reaching at least 10x the relative abundance of the blank sample in any of the experimental samples were classified as contaminants and removed [62].

#### Community composition analyses

Further analyses were performed and visualised using the ‘phyloseq’ package [63] with additional functions from ‘speedyseq’ [64] and ‘microbiome’ [65]. Visualisations were done with additional usage of ‘BioVenn’ [66, 67], ‘ggplot2’[68], ‘palletter’ [69], ‘RColorBrewer’ [70], ‘ggpubr’ [71], and ‘cowplot’ [72] packages. In data processing and analysis, packages: ‘Biostrings’ [73], ‘readxl’ [74], ‘writexl’ [75], ‘dplyr’ [76], and ‘tidyverse’ [77] were used.

α-diversity measures: the number of observed variants (ASV richness) and Shannon index were calculated on rarefied data using the estimate_richness function and visualized with the plot_richness function from ‘phyloseq’. Differences in α-diversity (ASV richness and Shannon index) between hive bees and caged bees, as well as between sampling timepoints (week 1 and week 2) and their interaction, were tested using a linear mixed-effects model with the lmer function from the packages ‘lme4’ [78] and ‘lmerTest’ [79]. DNA concentration in the sample and the number of individual samples in the pool were used as additional factors. Model performance was assessed using the ‘performance’ package [80]. To address autocorrelation and heteroscedasticity in residuals, the number of observed variants was log-transformed before modeling.

To test whether bacterial communities in bee samples and feeding sources differ, constrained correspondence analysis (CCA) with the ‘vegan’ package [81] was performed with sample type (bee sample vs. feeding source) as a variable. Further, β-diversity among bee samples was also analyzed using CCA. The model included bee source (hive vs. cage), timepoint (week 1 vs. week 2), and their interaction as predictors. Due to the scarcity of data and to improve model performance, we did not include replicates as factors in the analyses. A permutation test for the CCA model was performed using the anova.cca function from ‘vegan’ using 999 permutations. Divergence (heterogeneity of microbiota composition of samples from each bee source) was calculated as individual samples’ dissimilarities from respective group means using the Bray-Curtis dissimilarities with the divergence function in the ‘microbiome’ package. Differences in divergence between bee sources were tested with a linear model, the lm function in package ‘stats’ [51], after log-transformation with bee source, timepoint, and their interaction as predictors. To visualize the similarities between the microbial composition of bees and food/water samples, as well as between bee sources, non-metric multidimensional scaling (NMDS) plots based on Bray-Curtis dissimilarities were prepared using the plot_ordination function from the package ‘phyloseq’.

To visualise and perform tests on community composition at specific taxonomic levels, datasets were aggregated to the chosen taxonomic level using the aggregate_rare function, applying a detection cutoff of 300 reads per sample and a prevalence threshold of 20% of samples. Variants below these thresholds were categorized as “Other”. The microbial composition for the whole dataset was visualized on the family level and for the bee samples – additionally on the genus level. Heatmaps were generated on the bee samples data aggregated to the genus level.

Differential abundance of bacterial genera in the bee samples was analyzed using ‘DESeq2’ [82], with Benjamini-Hochberg correction for multiple comparisons [83], using the dataset aggregated to genus level as described above. All 20 genera resulting from the aggregation were included in the analysis. Genera abundances were considered statistically significant at p<0.05. Similarly, differential abundance of ASVs was analyzed using ‘DESeq2’ on count data. On the ASV level, the first two comparisons were conducted on ASV log-2-fold-change between hive bee crop vs. food samples and between hive bee crop vs. water samples, to identify variants significantly more abundant in the provided feeding sources than in the natural honeybee crop microbiota. These variants were marked as ‘bacteria enriched in food’ and ‘bacteria enriched in water’, respectively, in the ultimate ASVs comparison between hive bees vs. caged bees crop samples. In all the ASV differential abundance analyses, a significance threshold of p<0.01 was used to avoid discovering too many false-positive differences in ASV abundance.

## Results

### Ethanol content

Ethanol levels in the crop samples were consistently low across bee sources and timepoints, ranging from 0 to 0.35 g/L, with all group medians close to 0 (Fig. 1, Table 1). No ethanol was detected in water or food samples. There were no significant differences in ethanol levels between bee sources (χ^2^ = 0.2013, df = 1, p = 0.65), between weeks (χ^2^ = 0.3451, df = 1, p = 0.56), or in their interaction (χ^2^ = 0.9445, df = 1, p = 0.33). However, the zero-inflation component of the model indicated that caged bees had significantly more samples with detectable ethanol levels compared to hive bees (estimate ± SE = −1.09 ± 0.47, z = −2.291, p=0.022).

**Fig. 1.**
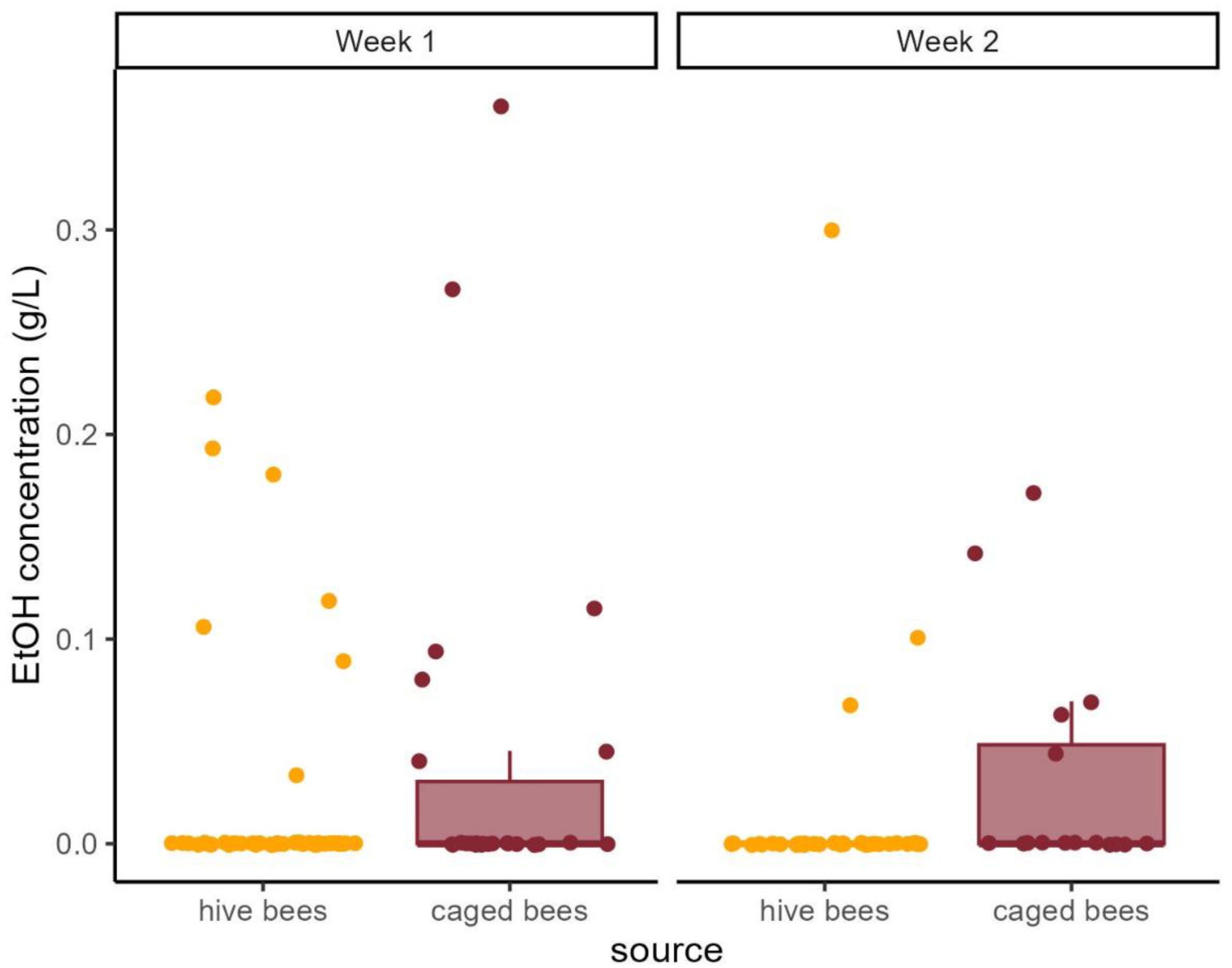
Ethanol concentration in crop samples, divided by bee source and timepoint (weeks). Dots indicate raw datapoints. Boxplots show medians and quartiles, with whiskers extending to 1.5 times the interquartile range.

**Table 1.** Ethanol level in crop samples – linear model results. The model was performed on log-transformed data. Intercept includes the following factor levels: source: hive bees and timepoint: week1.

| model | Estimate | Standard error | z value | Pr(>\|z\|) |
| --- | --- | --- | --- | --- |
| **Conditional model** |  |  |  |  |
| **intercept** | -2.00812 | 0.23125 | -8.684 | <2e-16 |
| **source: caged bees** | 0.06932 | 0.32704 | 0.212 | 0.832 |
| **timepoint: week2** | 0.15239 | 0.42221 | 0.361 | 0.718 |
| **Caged bees\*week2** | -0.53814 | 0.55372 | -0.972 | 0.331 |
| **Zero-inflation model** |  |  |  |  |
| **intercept** | 2.0149 | 0.3367 | 5.985 | 2.16e-09 |
| **Source: caged bees** | -1.0986 | 0.4796 | -2.291 | 0.022 |

### Microbiota

Sequencing of the libraries resulted in 22,264,863 read pairs, on average 473,720 per sample. Among them, after primer-trimming, dada2 pipeline identified 11,751 ASVs, out of which 7,255 were classified as bacterial ASVs. Among the bacterial ASVs, 151 were classified as contaminants based on their relative abundance in the blank sample. The distribution of contaminants across samples is shown in Online Resource 1: Fig. S1. The most common contaminants in the extraction blank belonged to the families: *Rhizobiaceae, Sphingomonadaceae,* and *Pseudomonadaceae* (Online Resource 1: Fig. S2), which is consistent with previous reports of common ultrapure water and Next Generation Sequencing contaminants [84, 85]. The final dataset included 7,104 ASVs, represented by 1– 1,826,393 read counts per variant.

#### α-diversity

A total of 875 ASVs were identified in caged bees and 2,431 ASVs in hive bees, with 234 ASVs (6.6%) shared between the two sources (Fig. 2a). ASV richness was significantly lower in caged bees than in hive bees (F=9.174, p=0.0076; Fig. 2b). However, neither timepoint (F=0.924, p=0.35), nor the interaction between bee source and timepoint (F=0.014, p=0.91) was significant. DNA concentration in the sample (F=1.338, p=0.26) as well as the number of samples in a pool (F=3.271, p=0.088) also had no significant effect on the ASV richness.

**Fig. 2.**
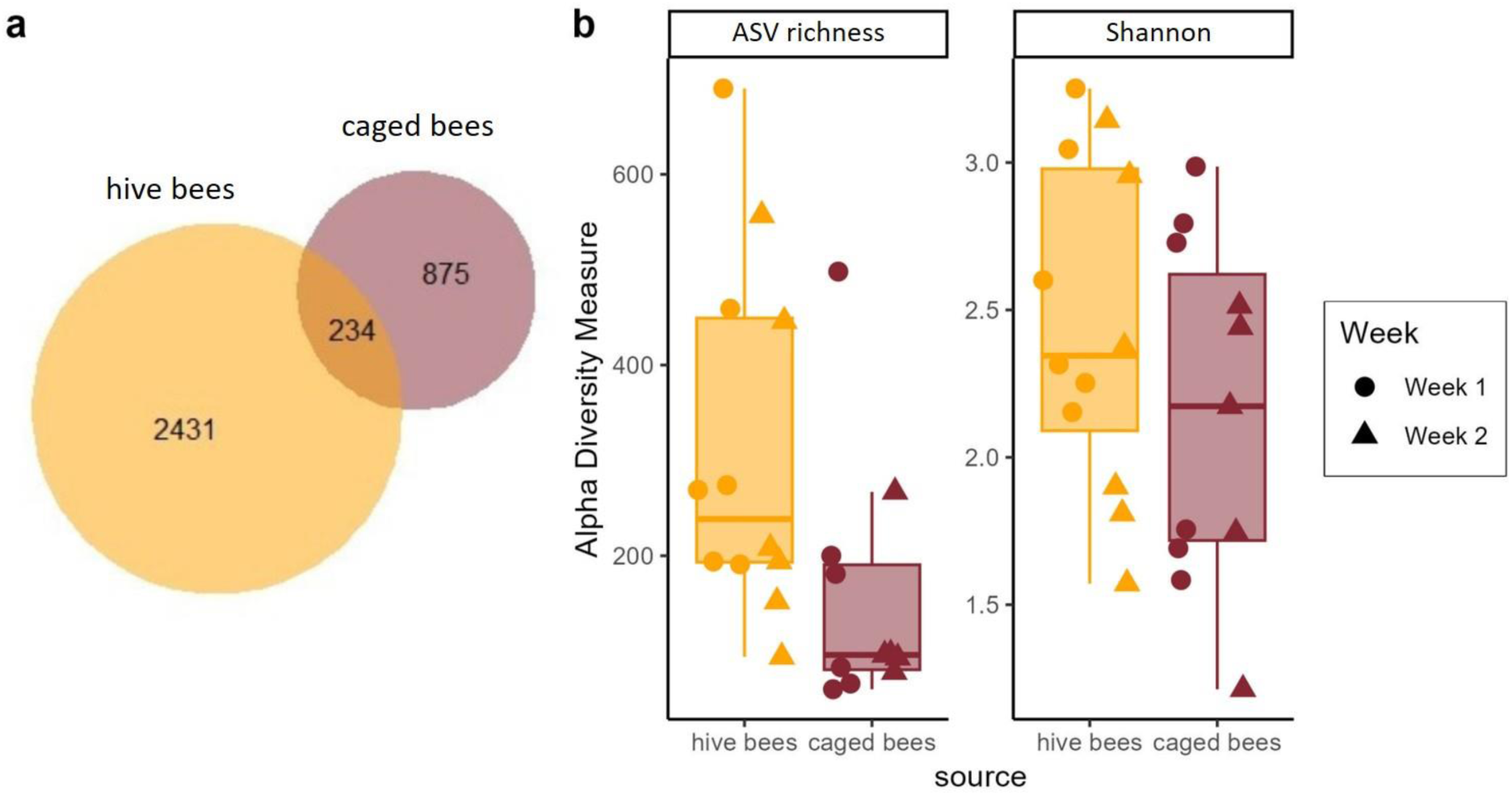
α-diversity measures. (a) Venn diagram showing the number of ASVs shared between hive and caged bees. (b) ASV richness (number of observed ASVs, left) and Shannon index (right) for hive and caged bees.

The Shannon index did not differ significantly between bee sources (F=1.485, p=0.24), timepoints (F=1.258, p=0.27), or their interaction (F=0.1371, p=0.72) (Fig. 2b). Neither DNA concentration in the sample (F=1.064, p=0.32) nor the number of samples in a pool (F=0.047, p=0.83) had a significant effect on the Shannon index.

#### β-diversity

Bacterial communities in samples from bees were different from those from feeding sources (CCA ANOVA: χ^2^=0.75, p=0.001, Fig. 3a). Bray-Curtis dissimilarity NMDS clustering revealed that food and water samples form distinct clusters, separated from bee samples (Fig. 3a), indicating that the microbial communities of bees and their feeding sources were compositionally distinct. Bee microbial communities from both sources (hive and caged) largely overlapped (Fig. 3a), though hive bees formed a more condensed cluster, while caged bees showed greater dispersion, with some samples clustering closely to hive bees and others overlapping with incubator food samples. Divergence was significantly higher in caged bees compared to hive bees: 0.265 (model ln estimate ± SE = −1.33 ± 0.07) for hive bees, 0.534 (−0.63 ± 0.07) for caged bees (Fig. 3b, Online Resource 1: Table S3).

**Fig. 3.**
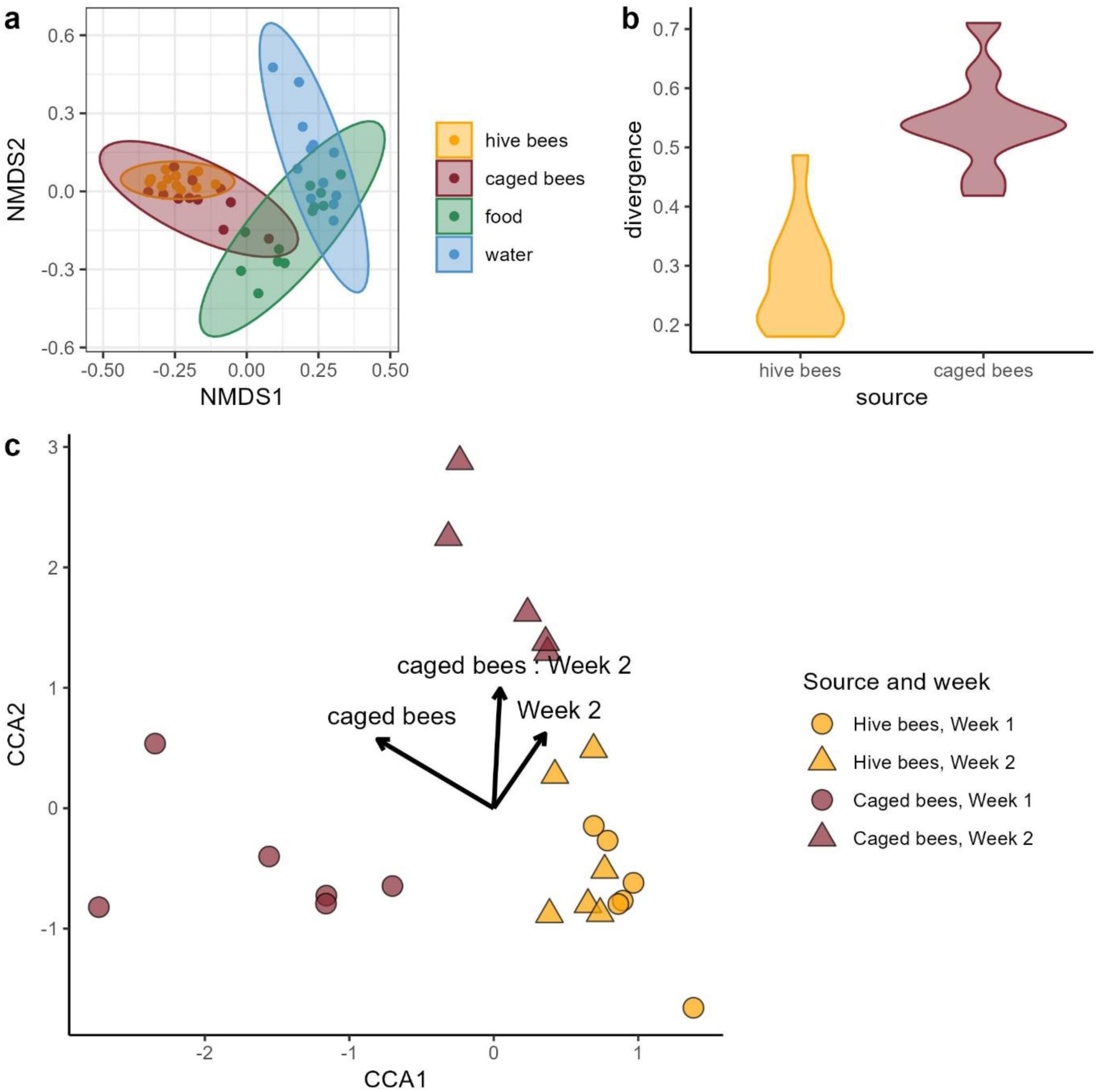
β-diversity plots. (a) Bray-Curtis distance NMDS ordination plots presenting clusters of hive bees, caged bees, food and water samples. Stress value = 0.15 (b) Violin plots showing divergence within each bee source. p<0.001. (c) CCA plot for the model including bee source, timepoint, and their interaction. CCA1 constrained axis explained 40% of the variance, CCA2 – 23%.

Constrained correspondence analysis (CCA) identified significant effects of the bee source and its interaction with timepoint on microbial community composition (Table 2, Fig. 3c). There was an evident separation between weeks in the caged bees (Fig. 3c). In total, the model explained 22.5% of the observed variance (Online Resource 1: Table S4).

**Table 2.** ANOVA test results for the CCA model. Df – degrees of freedom, Pr(>F) – p value.

| | Df | ChiSquare | F | $\text{Pr}( > F )$ |
| --- | --- | --- | --- | --- |
| **source** | 1 | 0.33679 | 2.5266 | 0.001 |
| **timepoint** | 1 | 0.18720 | 1.4044 | 0.074 |
| **source:timepoint** | 1 | 0.21144 | 1.5862 | 0.024 |
| **Residual** | 19 | 2.53271 |  |  |

#### Bacterial communities

At the family level, bee samples were mostly composed of *Lactobacillaceae* and *Acetobacteraceae*, which were almost absent from food and water samples (Fig. 4). Food and water samples exhibited greater diversity, with incubator food samples dominated by *Brevibacteriaceae*, which were also present in high abundance in two caged bee samples from week 1, but less abundant or absent in other caged bee or hive bee samples from week 2 (Fig. 4).

**Fig. 4.**
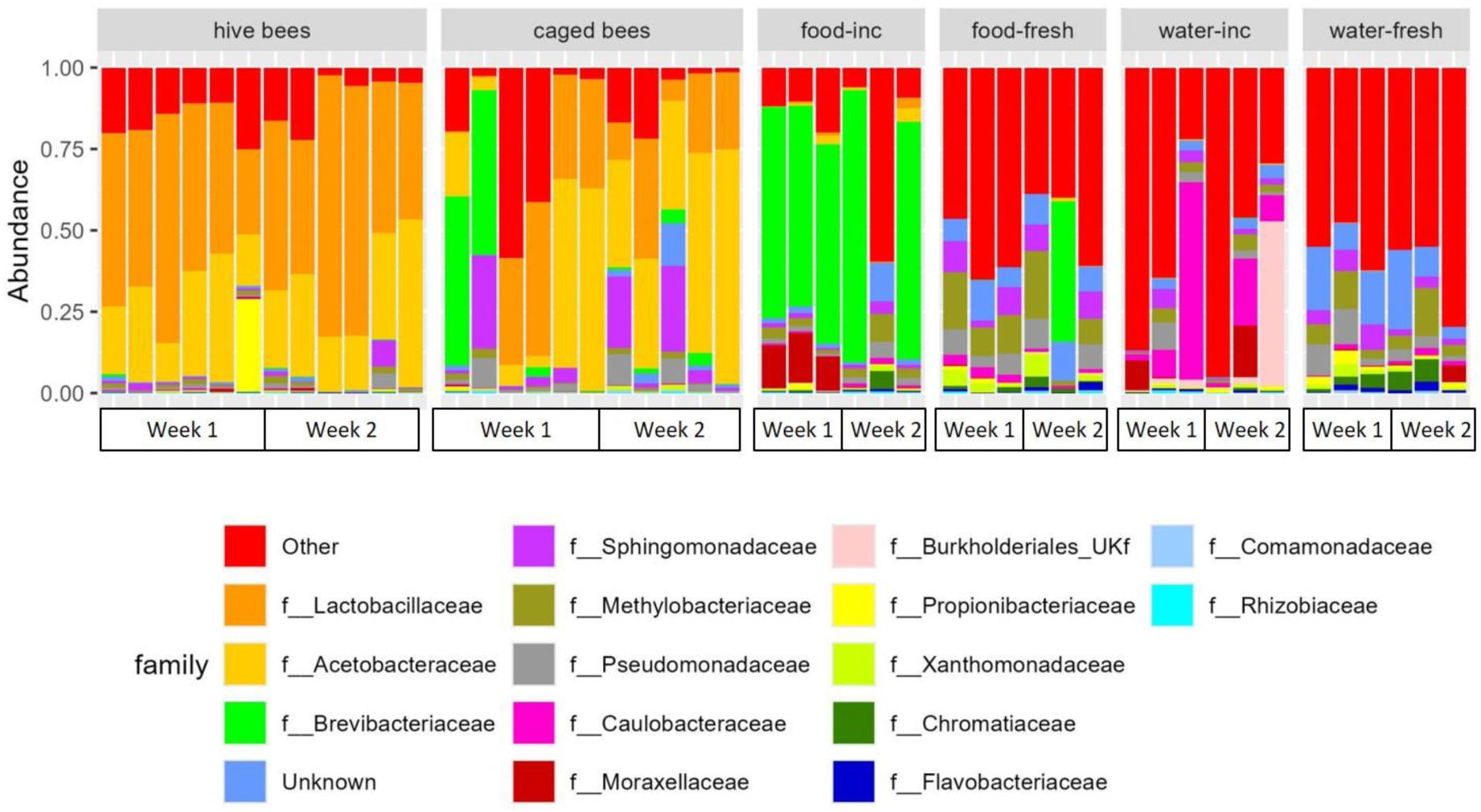
Bacterial communities composition on the family level in bee crop, food, and water samples. ‘inc’ – food and water samples after 24 h in the incubator, ‘fresh’ – freshly prepared.

The most abundant genera in both hive and caged bee samples were *Apilactobacillus* and *Bombella* (Fig. 5a, b). However, hive bees’ crop samples were characterized by a greater number of relatively abundant genera (Fig. 5c). The overall composition in hive bees was relatively similar between weeks, while the caged bees exhibited higher dissimilarity in week 1, becoming less heterogeneous in week 2 (Fig. 5a, c). On average, *Apilactobacillus* and *Bombella* together constituted over 75% of the crop bacterial community in hive bees, while in caged bees, these two dominant genera accounted for less than 50% of the crop microbiota (Fig. 5b). This difference was primarily driven by the higher abundance of *Apilactobacillus* in hive bees, while *Bombella* levels did not differ between bee sources (Fig. 5b, Fig. 6a). In contrast, caged bee samples were enriched in the next two common genera, *Sphingomonas* and *Pseudomonas* (Fig. 6a), and six less common genera: *Lactobacillus, Brevibacterium, Bifidobacterium, Snodgrassella, Gilliamella,* and *Delftia* (Fig. 6a). Genera enriched in the hive bees included, apart from the most common *Apilactobacillus,* also *Lonsdalea, Zymobacter, Acinetobacter, Fructobacillus,* and *Brevundimonas* (Fig. 6a). In total, out of the 20 compared genera, six were significantly less abundant in caged bees and eight were enriched in the caged bees (Fig. 6a).

**Fig. 5.**
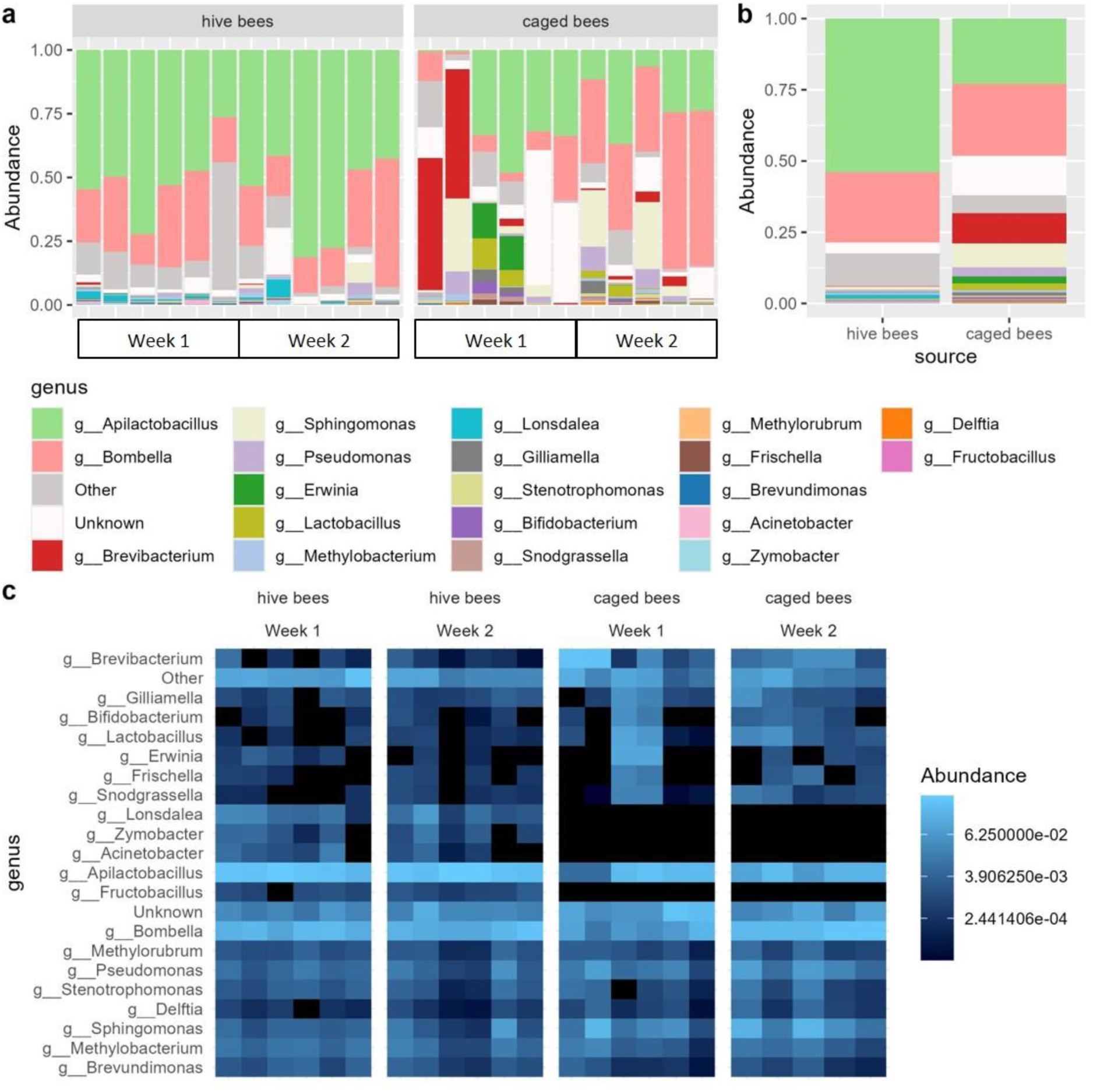
Bacterial communities composition on the genus level (a) in individual bee samples from hive bees (left) and caged bees (right); (b) in bee samples, averaged per bee source; (c) heatmaps divided by bee sources and weeks. Each column in (a) and (c) represents a single sample.

**Fig. 6.**
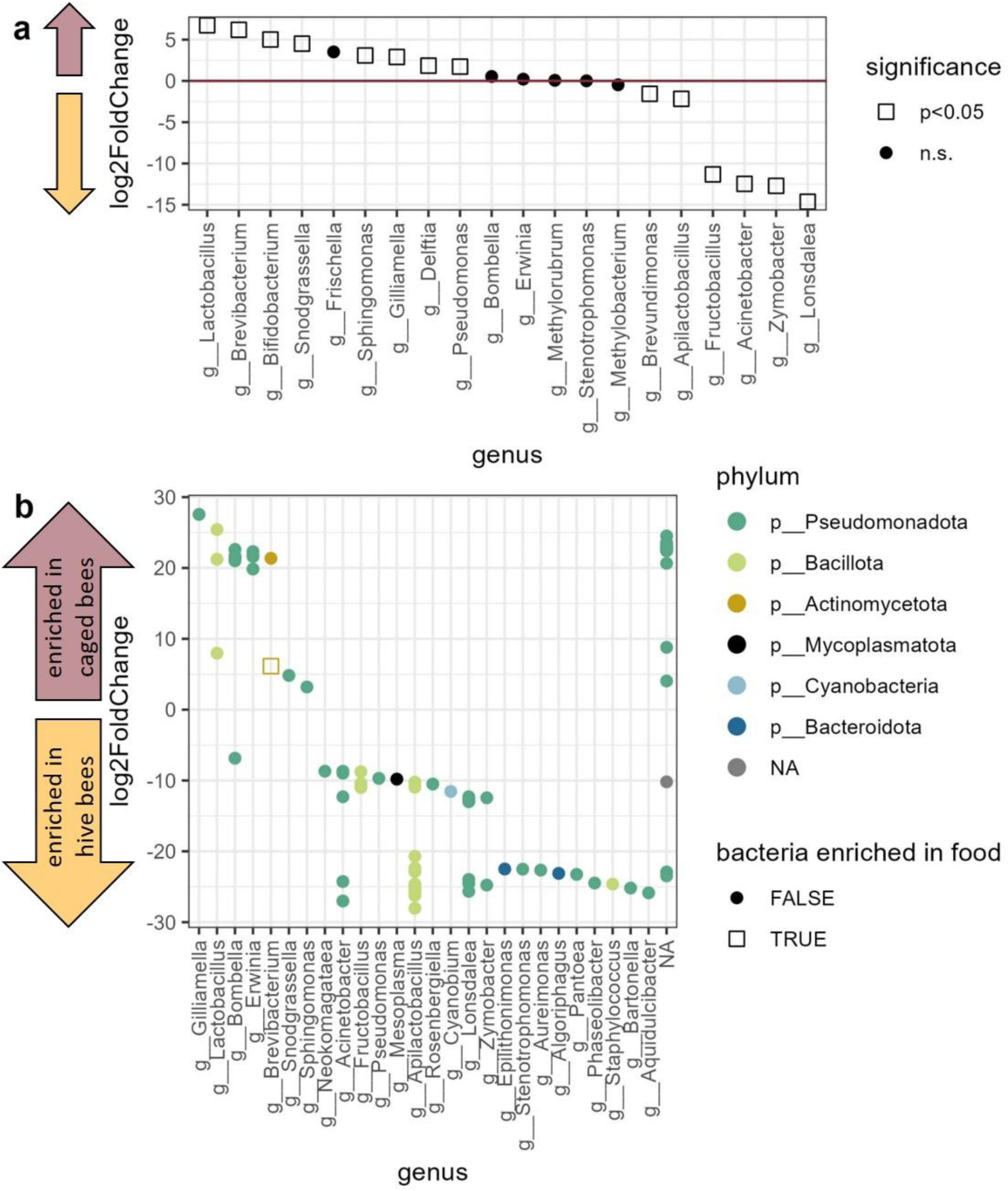
DESEq2 results. (a) Genera abundance log fold change on the bees dataset aggregated to the genus level. All genera included in the analysis are shown. Significance at the level p<0.05 (Benjamini-Hochberg corrected) is shown as point shapes; squares – significant, filled circles – not significant (n.s.). Genera more abundant in the caged bees are shown in the upper part of the plot (brown upward arrow), genera more abundant in the hive bees – in the lower part of the plot (golden downwards arrow) (b) ASVs log fold change between hive and caged bees. ASVs with positive log2fold change are enriched in caged bees (brown upward arrow) and with negative log2fold change – in hive bees (golden downwards arrow). Only significantly different (Benjamini-Hochberg corrected p<0.01) ASVs are shown.

In the differential abundance analysis (DESeq2) on ASV level, the comparison between hive bees and food samples identified 27 variants significantly more abundant in food than in the natural honeybee crop microbiota. These variants were classified as ‘bacteria enriched in food’ (Online Resource 1: Fig. S3, top panel). Analogously, 52 ASVs were identified as ‘bacteria enriched in water’ based on the comparison with water samples (Online Resource 1: Fig. S3, bottom panel). In the primary comparison between hive and caged bee samples, 57 ASVs were significantly more abundant in hive bees, while 34 ASVs were significantly more abundant in caged bees (p<0.01). Of the ASVs more abundant in caged bees, only one variant, belonging to *Brevibacterium,* was classified as ‘bacteria enriched in food’, while none were classified as ‘bacteria enriched in water’ (Fig. 6b). Among the core microbiota in the bee samples, two genera, *Apilactobacillus* and *Bombella,* showed contrasting patterns. Namely, *Apilactobacillus* had multiple variants overrepresented in hive bees and no ASVs characteristic for caged bees, while *Bombella* included several variants overrepresented in caged bees and only one variant that was significantly more abundant in hive bees. Other variants significantly more abundant in caged bees than in the hive bees were identified as representatives of a few other genera: *Gilliamella, Lactobacillus, Erwinia, Brevibacterium, Snodgrasella,* and *Sphingomonas*.

## Discussion

Our experiment showed that honeybees kept in laboratory conditions exhibit shifts in their crop microbiota compared to hive-reared bees. This effect was expected, as the crop microbiota is highly affected by the environmental supply of bacteria [47]. However, the crop microbiota of caged bees remained distinct from the microbial composition of food and water they received, but rather represented a reduced and more variable subset of the hive bee microbiota. This was observed as decreased ASV richness and higher β-diversity in caged bees compared to hive bees.

The most abundant bacterial genus in both groups, *Apilactobacillus*, was notably less abundant in caged bees, consistent with its known environmental origin. In contrast, *Bombella,* another core crop taxon, maintained stable abundance across groups. Other genera showed shifts in both directions. The increased relative abundance of *Gilliamella* and *Bifidobacterium* in caged bees raises the question of whether some taxa may contribute to endogenous ethanol accumulation, although this remains to be directly tested.

While ethanol content in the crop samples remained low (mean ∼0.01 g/L) and did not significantly differ in magnitude between groups, it was detected more frequently in caged bees. Some ethanol-positive samples were also observed in hive bees, consistent with the likelihood that bees encounter low levels of ethanol in the environment or within the hive [1, 17]. The detected levels were approximately two orders of magnitude higher than in haemolymph of bees reared under similar laboratory conditions [14] and in bees fed ethanol-spiked sucrose [86]. These levels are consistent with low-level ethanol presence in the crop and raise the possibility that fermentation may occur independently of environmental ethanol intake, although alternative explanations cannot be ruled out.

Despite a shared core composition (Fig. 3a), consistent with the core structure of the honeybee gut microbiota [33], the crop communities of caged bees were more heterogeneous (Fig. 3b), indicating potentially unstable microbial communities. This instability may reflect the unnatural, microbially impoverished incubator environment, which lacks the diverse microbial inputs typically encountered by foraging bees [28]. Hive microbiota is, on one hand, dependent on its surrounding environment, and on the other hand, serves as a microbial reservoir of high importance for colony health [87].

The low prevalence of bacteria dominant in caged bee samples within their food and water sources suggests minimal microbial transmission. In contrast, the temporary spike in *Brevibacterium* abundance observed in some caged bees in week 1 likely resulted from contamination with incubator food, where this genus was abundant (Fig. 5). This transient contamination appears to have destabilized the microbiota of some caged bees but was not observed in week 2, suggesting recovery and resilience. It also may have contributed to the greater differences observed between samples originating from caged bees, mainly driven by the samples from week 1 (Fig. 5).

Reduced ASV richness in caged bees compared to hive bees aligns with expectations, given the lack of environmental microbial inputs and the absence of hive-associated microbial exchange. This effect is consistent with previous findings that 22.5% of honeybee-associated bacterial variants, primarily those found on the body surface, are also present in flowers and are typically acquired indirectly through honey stored in the hive [47]. The two most abundant genera in both hive and caged bee samples, *Apilactobacillus* and *Bombella,* align with known crop microbiota components [18, 33–35, 40, 41, 88]. The most striking difference between hive and caged bees was observed in the reduced abundance of *Apilactobacillus* in caged bees (Fig. 5, Fig. 6a). This fructophilic genus, a known component of corbicular pollen [48] and a key player in floral microbiomes [35, 89], is likely acquired primarily from environmental sources. The absence of environmental sources in caged bees may account for their reduced *Apilactobacillus* levels.

Interestingly, *Bombella* abundance did not differ significantly between hive and caged bees. This stability may reflect a greater tolerance for laboratory diet caused by its preference for glucose as a carbon source [41], making this genus less sensitive to environmental isolation. In contrast, the genus *Gilliamella*, typically transmitted via social contact [46], especially with the fecal material [28], was more abundant in the crops of caged bees. This result is surprising, as previous studies have reported reduced *Gilliamella* abundance in the ileum of bees without hive contact [90], suggesting that the relationship between gut compartment and microbiota composition is complex and potentially environment-dependent.

*Lactobacillus* and *Bifidobacterium*, both of which are commonly found in the honeybee crop [28, 40, 49], were more abundant in caged bees than in hive bees. This increase might be plausibly explained by the stability and carbohydrate-rich nature of the laboratory diet, which favors lactic acid bacteria that can metabolize sucrose [91–93]. Similarly, *Acinetobacter,* one of the most abundant bacterial genera found in floral nectar [94], was rare in caged bees, likely reflecting the loss of regular environmental input. *Fructobacillus*, a fructose-specialist commonly found in flower nectar [95], was completely absent from the caged bees in this study. These observations are consistent with the idea that the loss of nectar-associated microbes in caged bees could create ecological opportunities for other taxa [40].

The hive bee crop microbiota was enriched in variants belonging to genera commonly associated with floral nectar, including *Acinetobacter, Fructobacillus, Pseudomonas, Rosenbergiella,* and *Pantoea* [94, 96]. Interestingly, one genus, *Bombella,* showed some strain-specificity, with separate variants enriched in caged and hive bees. This finding is in line with the generally reported high functional diversity within genera of bacterial microbionts [29–32, 97].

The presence or relative enrichment of taxa known to be capable of ethanol fermentation, such as *Gilliamella* and *Bifidobacterium*, although suggestive [33, 45, 46], does not confirm metabolic activity in vivo. Moreover, caging profoundly alters bee physiology and behavior, introducing multiple confounding factors. In such constrained conditions, they lack social cues and cannot forage outside, which increases their stress levels [98] and decreases immune responses [99]. Such changes may interfere with ethanol metabolism or clearance, potentially leading to low-level accumulation even without microbial fermentation. A more nuanced interpretation is therefore warranted: laboratory confinement may modulate both microbiota composition and host physiology, together enabling ethanol accumulation in the crop. To disentangle these mechanisms, future work should employ functionally explicit methods, such as ex vivo fermentation assays under crop-like conditions, metatranscriptomics or metabolomics to identify active fermentation pathways, or experiments with sterile bees and microbiota manipulations, e.g., antibiotics or gnotobiotic setups [29, 36, 37].

Our results are in line with recent findings [88] that the crop microbiota is shaped by the environmental sources only to some extent, much less than the mouth microbiota, and some ‘core taxa’ are present in the crop regardless of their external supply. This allows us to predict that some key microbiota functions are preserved even in the disturbed artificial environment. For example, abundances of several bacterial genera, known for their role in honeybee health [39, 40, 47], were not negatively affected (*Bombella)*, or even increased, like *Lactobacillus, Bifidobacterium, and Pseudomonas*. Also, bacteria involved in nutritional functions - carbohydrate utilization, as *Gilliamella* and *Snodgrassella,* which engage in cross-feeding [37, 100], were highly abundant in caged bees. However, it should be noted that as relative abundances were analyzed, it cannot be determined if particular bacteria were less or more abundant in terms of absolute values between bee sources.

## Conclusions

Our findings are consistent with the hypothesis that laboratory confinement reduces crop microbiota richness in honeybees and is associated with increased relative abundance of opportunistic and potentially ethanol-producing taxa such as *Gilliamella* and *Bifidobacterium*. At the same time, some core components, like *Bombella*, remained stable, suggesting the resilience of key symbionts. However, the precise relationship between these microbial shifts and ethanol production remains to be fully explored. Future studies should also investigate how ethanol exposure itself may shape the microbiota and should integrate both bacterial and fungal communities, especially given the influential roles of fungi in crop microbiota [101] (but see [102]). Assessing the functional consequences of these shifts, including potential impacts on host physiology, behavior, and health, is essential for understanding how microbial symbionts contribute to honeybee phenotypic plasticity.

## Supporting information

Online Resource 1

## Statements and declarations

### Author contributions

Conceptualization: Krzysztof Miler; Formal analysis: Weronika Antoł, Bartłomiej Surmacz, Krzysztof Miler; Funding acquisition: Krzysztof Miler; Investigation: Weronika Antoł, Krzysztof Miler, Monika Ostap-Chec, Daniel Stec; Methodology: all authors; Resources: Monika Ostap-Chec, Daniel Stec, Krzysztof Miler; Visualization: Weronika Antoł, Bartłomiej Surmacz; Writing – original draft: Weronika Antoł, Krzysztof Miler; Writing – review & editing: all authors.

### Ethics approval

No approval of research ethics committees was required to accomplish the goals of this study because experimental work was conducted with an unregulated invertebrate species.

## Literature

1. Bowland AC, Melin AD, Hosken DJ, et al (2025) The evolutionary ecology of ethanol. Trends Ecol Evol 40:67–79. 10.1016/j.tree.2024.09.005

2. Madden AA, Epps MJ, Fukami T, et al (2018) The ecology of insect–yeast relationships and its relevance to human industry. Proc R Soc B Biol Sci 285:20172733. 10.1098/rspb.2017.2733

3. Montgomery ME, Wargo PM (1983) Ethanol and other host-derived volatiles as attractants to beetles that bore into hardwoods. J Chem Ecol 9:181–190. 10.1007/BF00988035

4. Nout MJR, Bartelt RJ (1998) Attraction of a flying nitulid *Carpophilus humeralis* to volatiles produced by yeasts grown on sweet corn and corn-based medium. Journal of Chemical Ecology 24:1217–1239

5. Beaulieu M, Franke K, Fischer K (2017) Feeding on ripening and over-ripening fruit: interactions between sugar, ethanol and polyphenol contents in a tropical butterfly. J Exp Biol 220:3127–3134. 10.1242/jeb.162008

6. Azanchi R, Kaun KR, Heberlein U (2013) Competing dopamine neurons drive oviposition choice for ethanol in Drosophila. Proc Natl Acad Sci U S A 110:21153– 21158. 10.1073/pnas.1320208110

7. Sokolowski MBC, Abramson CI, Craig DPA (2012) Ethanol Self-Administration in Free-Flying Honeybees ( *Apis mellifera* L.) in an Operant Conditioning Protocol. Alcohol Clin Exp Res 36:1568–1577. 10.1111/j.1530-0277.2012.01770.x

8. Mustard JA, Oquita R, Garza P, Stoker A (2019) Honey Bees (*Apis mellifera*) Show a Preference for the Consumption of Ethanol. Alcohol Clin Exp Res 43:26–35. 10.1111/acer.13908

9. Bouchebti S, Gershon Y, Gordin A, et al (2024) Tolerance and efficient metabolization of extremely high ethanol concentrations by a social wasp. Proc Natl Acad Sci 121:e2410874121. 10.1073/pnas.2410874121

10. Milan NF, Kacsoh BZ, Schlenke TA (2012) Alcohol consumption as self-medication against blood-borne parasites in the fruit fly. Curr Biol CB 22:488–493. 10.1016/j.cub.2012.01.045

11. Castillo C, Chen H, Graves C, et al (2012) Biosynthesis of ethyl oleate, a primer pheromone, in the honey bee (Apis mellifera L.). Insect Biochem Mol Biol 42:404–416. 10.1016/j.ibmb.2012.02.002

12. Leoncini I, Le Conte Y, Costagliola G, et al (2004) Regulation of behavioral maturation by a primer pheromone produced by adult worker honey bees. Proc Natl Acad Sci 101:17559–17564. 10.1073/pnas.0407652101

13. Miler K, Stec D, Kamińska A, et al (2021) Alcohol intoxication resistance and alcohol dehydrogenase levels differ between the honeybee castes. Apidologie 52:230–241. 10.1007/s13592-020-00812-y

14. Ostap-Chec M, Antoł W, Bajorek D, et al (2025) Honeybees show an increased preference for dietary alcohol when parasitized

15. Ostap-Chec M, Bajorek D, Antoł W, et al (2024) Occasional and constant exposure to dietary ethanol shortens the lifespan of worker honey bees. J Comp Physiol B 194:403–410. 10.1007/s00360-024-01571-3

16. Ostap-Chec M, Bajorek D, Antoł W, et al (2025) Honey bees are resilient to the long-term presence of alcohol in their diet. Ecotoxicology. 10.1007/s10646-025-02903-x

17. Miler K (2025) Ethanol and pollinators: expanding Bowland et al.’s framework. Trends Ecol Evol 40:115–116. 10.1016/j.tree.2024.11.019

18. Johnson BR (2023) Honey bee biology, First edition. Princeton University Press, Princeton Oxford

19. Crailsheim K (1988) Regulation of food passage in the intestine of the honeybee (Apis mellifera L.). Insect Physiol 34:85–90. 10.1016/0022-1910(88)90158-8

20. Carreck NL, Andree M, Brent CS, et al (2013) Standard methods for *Apis mellifera* anatomy and dissection. J Apic Res 52:1–40. 10.3896/IBRA.1.52.4.03

21. Bayoumy AB, Mulder CJJ, Mol JJ, Tushuizen ME (2021) Gut fermentation syndrome: A systematic review of case reports. United Eur Gastroenterol J 9:332–342. 10.1002/ueg2.12062

22. Kruckenberg KM, DiMartini AF, Rymer JA, et al (2020) Urinary Auto-brewery Syndrome: A Case Report. Ann Intern Med 172:702–704. 10.7326/L19-0661

23. Tamama K, Kruckenberg KM, DiMartini AF (2024) Gut and bladder fermentation syndromes: a narrative review. BMC Med 22:26. 10.1186/s12916-023-03241-7

24. Elshaghabee FMF, Bockelmann W, Meske D, et al (2016) Ethanol Production by Selected Intestinal Microorganisms and Lactic Acid Bacteria Growing under Different Nutritional Conditions. Front Microbiol 7:. 10.3389/fmicb.2016.00047

25. Dinis-Oliveira RJ (2021) The Auto-Brewery Syndrome: A Perfect Metabolic “Storm” with Clinical and Forensic Implications. J Clin Med 10:4637. 10.3390/jcm10204637

26. Paramsothy J, Gutlapalli SD, Ganipineni VDP, et al (2023) Understanding Auto-Brewery Syndrome in 2023: A Clinical and Comprehensive Review of a Rare Medical Condition. Cureus 15:. 10.7759/cureus.37678

27. Šoša I (2023) The Human Body as an Ethanol-Producing Bioreactor—The Forensic Impacts. Fermentation 9:738. 10.3390/fermentation9080738

28. Powell JE, Martinson VG, Urban-Mead K, Moran NA (2014) Routes of Acquisition of the Gut Microbiota of the Honey Bee Apis mellifera. Appl Environ Microbiol 80:7378–7387. 10.1128/AEM.01861-14

29. Engel P, Martinson VG, Moran NA (2012) Functional diversity within the simple gut microbiota of the honey bee. Proc Natl Acad Sci 109:11002–11007. 10.1073/pnas.1202970109

30. Ellegaard KM, Tamarit D, Javelind E, et al (2015) Extensive intra-phylotype diversity in lactobacilli and bifidobacteria from the honeybee gut. BMC Genomics 16:284. 10.1186/s12864-015-1476-6

31. Ellegaard KM, Engel P (2019) Genomic diversity landscape of the honey bee gut microbiota. Nat Commun 10:446. 10.1038/s41467-019-08303-0

32. Yang C, Hu J, Su Q, et al (2025) A review on recent taxonomic updates of gut bacteria associated with social bees, with a curated genomic reference database. Insect Sci 32:2–23. 10.1111/1744-7917.13365

33. Kwong WK, Moran NA (2016) Gut microbial communities of social bees. Nat Rev Microbiol 14:374–384. 10.1038/nrmicro.2016.43

34. Smith EA, Anderson KE, Corby-Harris V, et al (2020) Reclassification of seven honey bee symbiont strains as *Bombella apis*

35. Zheng J, Wittouck S, Salvetti E, et al (2020) A taxonomic note on the genus Lactobacillus: Description of 23 novel genera, emended description of the genus Lactobacillus Beijerinck 1901, and union of Lactobacillaceae and Leuconostocaceae. Int J Syst Evol Microbiol 70:2782–2858. 10.1099/ijsem.0.004107

36. Lee FJ, Rusch DB, Stewart FJ, et al (2015) Saccharide breakdown and fermentation by the honey bee gut microbiome. Environ Microbiol 17:796–815. 10.1111/1462-2920.12526

37. Kešnerová L, Mars RAT, Ellegaard KM, et al (2017) Disentangling metabolic functions of bacteria in the honey bee gut. PLOS Biol 15:e2003467. 10.1371/journal.pbio.2003467

38. Kešnerová L, Emery O, Troilo M, et al (2020) Gut microbiota structure differs between honeybees in winter and summer. ISME J 14:801–814. 10.1038/s41396-019-0568-8

39. Forsgren E, Olofsson TC, Vásquez A, Fries I (2010) Novel lactic acid bacteria inhibiting *Paenibacillus larvae* in honey bee larvae. Apidologie 41:99–108. 10.1051/apido/2009065

40. Vásquez A, Forsgren E, Fries I, et al (2012) Symbionts as Major Modulators of Insect Health: Lactic Acid Bacteria and Honeybees. PLoS ONE 7:e33188. 10.1371/journal.pone.0033188

41. Anderson KE, Copeland DC (2024) The honey bee “hive” microbiota: meta-analysis reveals a native and aerobic microbiota prevalent throughout the social resource niche. Front Bee Sci 2:1410331. 10.3389/frbee.2024.1410331

42. El Khoury S, Gauthier J, Mercier PL, et al (2024) Honeybee gut bacterial strain improved survival and gut microbiota homeostasis in *Apis mellifera* exposed *in vivo* to clothianidin. Microbiol Spectr 12:e00578–24. 10.1128/spectrum.00578-24

43. Liberti J, Kay T, Quinn A, et al (2022) The gut microbiota affects the social network of honeybees. Nat Ecol Evol 6:1471–1479. 10.1038/s41559-022-01840-w

44. Cabirol A, Moriano-Gutierrez S, Engel P (2024) Neuroactive metabolites modulated by the gut microbiota in honey bees. Mol Microbiol 122:284–293. 10.1111/mmi.15167

45. Van der Meulen R, Adriany T, Verbrugghe K, De Vuyst L (2006) Kinetic Analysis of Bifidobacterial Metabolism Reveals a Minor Role for Succinic Acid in the Regeneration of NAD+ through Its Growth-Associated Production. Appl Environ Microbiol 72:5204–5210. 10.1128/AEM.00146-06

46. Nguyen VH (2024) Genomic investigations of diverse corbiculate bee gut-associated Gilliamella reveal conserved pathways for energy metabolism, with diverse and variable energy sources. Access Microbiol 6:. 10.1099/acmi.0.000793.v3

47. Tiusanen M, Becker-Scarpitta A, Wirta H (2024) Distinct Communities and Differing Dispersal Routes in Bacteria and Fungi of Honey Bees, Honey, and Flowers. Microb Ecol 87:100. 10.1007/s00248-024-02413-z

48. Graystock P, Rehan SM, McFrederick QS (2017) Hunting for healthy microbiomes: determining the core microbiomes of Ceratina, Megalopta, and Apis bees and how they associate with microbes in bee collected pollen. Conserv Genet 18:701–711. 10.1007/s10592-017-0937-7

49. Olofsson TC, Vásquez A (2008) Detection and Identification of a Novel Lactic Acid Bacterial Flora Within the Honey Stomach of the Honeybee Apis mellifera. Curr Microbiol 57:356–363. 10.1007/s00284-008-9202-0

50. Vásquez A, Olofsson TC (2009) The lactic acid bacteria involved in the production of bee pollen and bee bread. J Apic Res 48:189–195. 10.3896/IBRA.1.48.3.07

51. R Core Team (2024) R: A Language and Environment for Statistical Computing

52. Brooks ME, Kristensen K, Benthem KJ van, et al (2017) glmmTMB Balances Speed and Flexibility Among Packages for Zero-inflated Generalized Linear Mixed Modeling. The R Journal 9:378–400. 10.32614/RJ-2017-066

53. Hartig F (2024) DHARMa: Residual Diagnostics for Hierarchical (Multi-Level / Mixed) Regression Models

54. Illumina (2013) 16S Metagenomic Sequencing Library Preparation. https://support.illumina.com/documents/documentation/chemistry_documentation/16s/16s-metagenomic-library-prep-guide-15044223-b.pdf#page=20.14. Accessed 19 May 2025

55. Takahashi S, Tomita J, Nishioka K, et al (2014) Development of a Prokaryotic Universal Primer for Simultaneous Analysis of Bacteria and Archaea Using Next-Generation Sequencing. PLoS ONE 9:e105592. 10.1371/journal.pone.0105592

56. Martin M (2011) Cutadapt removes adapter sequences from high-throughput sequencing reads. EMBnet.journal 17:10–12. 10.14806/ej.17.1.200

57. Callahan BJ, McMurdie PJ, Rosen MJ, et al (2016) DADA2: High-resolution sample inference from Illumina amplicon data. Nat Methods 13:581–583. 10.1038/nmeth.3869

58. Callahan B (2016) DADA2 tutorial. https://benjjneb.github.io/dada2/tutorial.html

59. Murali A, Bhargava A, Wright ES (2018) IDTAXA: a novel approach for accurate taxonomic classification of microbiome sequences. Microbiome 6:140. 10.1186/s40168-018-0521-5

60. Wright E S (2016) Using DECIPHER v2.0 to Analyze Big Biological Sequence Data in R. R J 8:352. 10.32614/RJ-2016-025

61. Daisley BA, Reid G (2021) BEExact: a Metataxonomic Database Tool for High-Resolution Inference of Bee-Associated Microbial Communities. mSystems 6:10.1128/msystems.00082-21. 10.1128/msystems.00082-21

62. Łukasik P, Newton JA, Sanders JG, et al (2017) The structured diversity of specialized gut symbionts of the New World army ants. Mol Ecol 26:3808–3825. 10.1111/mec.14140

63. McMurdie PJ, Holmes S (2013) phyloseq: An R Package for Reproducible Interactive Analysis and Graphics of Microbiome Census Data. PLoS ONE 8:e61217. 10.1371/journal.pone.0061217

64. McLaren M (2025) speedyseq: Faster implementations of phyloseq functions

65. Lahti L, Shetty S (2012) microbiome R package

66. Hulsen T (2021) BioVenn: Create Area-Proportional Venn Diagrams from Biological Lists

67. Hulsen T (2021) BioVenn – an R and Python package for the comparison and visualization of biological lists using area-proportional Venn diagrams. Data Science 4:51–61

68. Wickham H (2016) ggplot2: Elegant Graphics for Data Analysis

69. Hvitfeldt E (2025) paletteer: Comprehensive Collection of Color Palettes

70. Neuwirth E (2022) RColorBrewer: ColorBrewer Palettes

71. Kassambara A (2023) ggpubr: “ggplot2” Based Publication Ready Plots

72. Wilke C (2024) cowplot: Streamlined Plot Theme and Plot Annotations for ‘ggplot2

73. Pagès H, Aboyoun P, Gentleman R, DebRoy S (2024) Biostrings: Efficient manipulation of biological strings

74. Wickham H, Bryan J (2023) readxl: Read Excel Files

75. Ooms J (2024) writexl: Export Data Frames to Excel “xlsx” Format

76. Wickham H, François R, Henry L, et al (2023) dplyr: A Grammar of Data Manipulation

77. Wickham H, Averick M, Bryan J, et al (2019) Welcome to the Tidyverse. J Open Source Softw 4:1686. 10.21105/joss.01686

78. Bates D, Mächler M, Bolker B, Walker S (2015) Fitting Linear Mixed-Effects Models Using **lme4**. J Stat Softw 67:. 10.18637/jss.v067.i01

79. Kuznetsova A, Brockhoff PB, Christensen RHB (2017) **lmerTest** Package: Tests in Linear Mixed Effects Models. J Stat Softw 82:. 10.18637/jss.v082.i13

80. Lüdecke D, Ben-Shachar M, Patil I, et al (2021) performance: An R Package for Assessment, Comparison and Testing of Statistical Models. J Open Source Softw 6:3139. 10.21105/joss.03139

81. Oksanen J, Simpson GL, Blanchet FG, et al (2025) vegan: Community Ecology Package

82. Love MI, Huber W, Anders S (2014) Moderated estimation of fold change and dispersion for RNA-seq data with DESeq2. Genome Biol 15:550. 10.1186/s13059-014-0550-8

83. Benjamini Y, Hochberg Y (1995) Controlling the False Discovery Rate: A Practical and Powerful Approach to Multiple Testing. J R Stat Soc Ser B Stat Methodol 57:289–300. 10.1111/j.2517-6161.1995.tb02031.x

84. Kéki Z, Grébner K, Bohus V, et al (2013) Application of special oligotrophic media for cultivation of bacterial communities originated from ultrapure water. Acta Microbiol Immunol Hung 60:345–357. 10.1556/AMicr.60.2013.3.9

85. Laurence M, Hatzis C, Brash DE (2014) Common Contaminants in Next-Generation Sequencing That Hinder Discovery of Low-Abundance Microbes. PLoS ONE 9:e97876. 10.1371/journal.pone.0097876

86. Golańska M, Antoł W, Witek M, Miler K (2025) Buzzed but not elated? Effect of ethanol on cognitive judgement bias in honeybees. BioRxiv

87. Gorrochategui-Ortega J, Muñoz-Colmenero M, Kovačić M, et al (2022) A short exposure to a semi-natural habitat alleviates the honey bee hive microbial imbalance caused by agricultural stress. Sci Rep 12:18832. 10.1038/s41598-022-23287-6

88. Warren ML, Tsuji K, Decker LE, et al (2025) Bacteria in honeybee crops are decoupled from those in floral nectar and bee mouths. 88:. 10.1007/s00248-025-02544-x

89. Maeno S, Tanizawa Y, Kanesaki Y, et al (2016) Genomic characterization of a fructophilic bee symbiont Lactobacillus kunkeei reveals its niche-specific adaptation. Syst Appl Microbiol 39:516–526. 10.1016/j.syapm.2016.09.006

90. Anderson KE, Ricigliano VA, Copeland DC, et al (2023) Social Interaction is Unnecessary for Hindgut Microbiome Transmission in Honey Bees: The Effect of Diet and Social Exposure on Tissue-Specific Microbiome Assembly. Microb Ecol 85:1498–1513. 10.1007/s00248-022-02025-5

91. Gänzle MG, Follador R (2012) Metabolism of Oligosaccharides and Starch in Lactobacilli: A Review. Front Microbiol 3:. 10.3389/fmicb.2012.00340

92. Levantovsky R, Allen-Blevins C, Sela D (2018) Nutritional Requirements of Bifidobacteria. In: The Bifidobacteria and Related Organisms. pp 115–129

93. Gouda MNR, Subramanian S, Kumar A, Ramakrishnan B (2024) Microbial ensemble in the hives: deciphering the intricate gut ecosystem of hive and forager bees of Apis mellifera. Mol Biol Rep 51:262. 10.1007/s11033-024-09239-5

94. Quevedo-Caraballo S, De Vega C, Lievens B, et al (2025) Tiny but mighty? Overview of a decade of research on nectar bacteria. New Phytol 245:1897–1910. 10.1111/nph.20369

95. Endo A, Maeno S, Tanizawa Y, et al (2018) Fructophilic Lactic Acid Bacteria, a Unique Group of Fructose-Fermenting Microbes. Appl Environ Microbiol 84:e01290–18. 10.1128/AEM.01290-18

96. Morris MM, Frixione NJ, Burkert AC, et al (2020) Microbial abundance, composition, and function in nectar are shaped by flower visitor identity. FEMS Microbiol Ecol 96:fiaa003. 10.1093/femsec/fiaa003

97. Ludvigsen J, Porcellato D, Amdam GV, Rudi K (2018) Addressing the diversity of the honeybee gut symbiont Gilliamella: description of Gilliamella apis sp. nov., isolated from the gut of honeybees (Apis mellifera). Int J Syst Evol Microbiol 68:1762–1770. 10.1099/ijsem.0.002749

98. Lattorff HMG (2022) Increased Stress Levels in Caged Honeybee (Apis mellifera) (Hymenoptera: Apidae) Workers. Stresses 2:373–383. 10.3390/stresses2040026

99. Gregory CL, Fell RD, Belden LK, Walke JB (2022) Classic Hoarding Cages Increase Gut Bacterial Abundance and Reduce the Individual Immune Response of Honey Bee (Apis mellifera) Workers. J Insect Sci 22:6. 10.1093/jisesa/ieac016

100. Kwong WK, Engel P, Koch H, Moran NA (2014) Genomics and host specialization of honey bee and bumble bee gut symbionts. Proc Natl Acad Sci 111:11509–11514. 10.1073/pnas.1405838111

101. Callegari M, Crotti E, Fusi M, et al (2021) Compartmentalization of bacterial and fungal microbiomes in the gut of adult honeybees. Npj Biofilms Microbiomes 7:42. 10.1038/s41522-021-00212-9

102. Decker LE, San Juan PA, Warren ML, et al (2023) Higher Variability in Fungi Compared to Bacteria in the Foraging Honey Bee Gut. Microb Ecol 85:330–334. 10.1007/s00248-021-01922-5

