## Supplementary material for "Do shifts in honeybee crop microbiota enable ethanol accumulation? A comparative analysis of caged and foraging bees": Online Resource 1

Table S1. Number of sample pools per category for ethanol level measurement and microbiota analysis.

| | **Ethanol** | | **Microbiota** | |
|---|---|---|---|---|
| **Sample source** | Week 1 | Week 2 | Week 1 | Week 2 |
| **Bee crop** | | | | |
| caged bees | 26 | 16 | 6 | 5 |
| hive bees | 48 | 37 | 6 | 6 |
| **Food** | | | | |
| fresh | 6 | 6 | 3 | 3 |
| incubator | 6 | 6 | 3 | 3 |
| **Water** | | | | |
| fresh | 6 | 6 | 3 | 3 |
| incubator | 6 | 6 | 3 | 3 |

Table S2. Number of reads for each sample at several stages of data preprocessing: original raw reads (INPUT), after: quality filtering and trimming (FILTER), denoising (DENOISED_F and DENOISED_R for forward and reversed reads, respectively), merging the F and R reads (MERGED), removing chimeras (NONCHIM). The last column (%) shows the percentage of original reads passed to further analysis.

| | INPUT | FILTERED | DENOISED_F | DENOISED_R | MERGED | NONCHIM | % |
|---|---|---|---|---|---|---|---|
| **M1-R2-T1-C-1-16S** | 540343 | 446835 | 444306 | 444492 | 392587 | 386086 | 71% |
| **M1-R2-T1-C-2-16S** | 434538 | 354571 | 353479 | 353375 | 337318 | 329287 | 76% |
| **M1-R2-T1-U-1-16S** | 718100 | 616470 | 613101 | 611679 | 460375 | 446688 | 62% |
| **M1-R2-T1-U-2-16S** | 278047 | 225292 | 224157 | 224002 | 168204 | 164261 | 59% |
| **M1-R2-T1-P-C-16S** | 312155 | 248430 | 247666 | 247499 | 239338 | 238043 | 76% |
| **M1-R2-T1-P-F-16S** | 568769 | 482788 | 481417 | 482187 | 404857 | 401797 | 71% |
| **M1-R2-T1-W-C-16S** | 592663 | 493253 | 491085 | 491555 | 454457 | 446904 | 75% |
| **M1-R2-T1-W-F-16S** | 402936 | 332853 | 326497 | 326360 | 206103 | 186909 | 46% |
| **M1-R2-T2-C-1-16S** | 246322 | 215608 | 214811 | 214970 | 175048 | 166742 | 68% |
| **M1-R2-T2-C-2-16S** | 1326357 | 1097069 | 1093191 | 1093653 | 832342 | 798789 | 60% |
| **M1-R2-T2-U-1-16S** | 657790 | 538773 | 535814 | 535499 | 398665 | 389369 | 59% |
| **M1-R2-T2-U-2-16S** | 614716 | 485976 | 483158 | 483070 | 373299 | 363301 | 59% |
| **M1-R2-T2-P-C-16S** | 320494 | 233639 | 231176 | 233198 | 226439 | 225646 | 70% |
| **M1-R2-T2-P-F-16S** | 463522 | 379996 | 378239 | 379138 | 335621 | 327359 | 71% |
| **M1-R2-T2-W-C-16S** | 445788 | 345125 | 341865 | 342896 | 233611 | 232285 | 52% |
| **M1-R2-T2-W-F-16S** | 167222 | 144052 | 141881 | 142027 | 104903 | 103762 | 62% |
| **M1-R3-T1-C-1-16S** | 312758 | 190533 | 189598 | 189896 | 137226 | 131271 | 42% |
| **M1-R3-T1-C-2-16S** | 970664 | 790233 | 788232 | 787627 | 567553 | 542254 | 56% |
| **M1-R3-T1-U-1-16S** | 258843 | 207918 | 206916 | 207057 | 163422 | 160840 | 62% |

| | | | | | | | |
|---|---|---|---|---|---|---|---|
| **M1-R3-T1-U-2-16S** | 363985 | 246672 | 246012 | 245346 | 188348 | 186443 | 51% |
| **M1-R3-T1-P-C-16S** | 273822 | 222995 | 222050 | 222578 | 212140 | 211607 | 77% |
| **M1-R3-T1-P-F-16S** | 96262 | 45382 | 44888 | 45091 | 39893 | 39840 | 41% |
| **M1-R3-T1-W-C-16S** | 255147 | 202455 | 201319 | 201819 | 170404 | 169035 | 66% |
| **M1-R3-T1-W-F-16S** | 517119 | 432405 | 429806 | 431440 | 384516 | 379034 | 73% |
| **M1-R3-T2-C-1-16S** | 581936 | 493501 | 492383 | 492708 | 389446 | 368306 | 63% |
| **M1-R3-T2-C-2-16S** | 425361 | 353832 | 353113 | 353240 | 271733 | 264471 | 62% |
| **M1-R3-T2-U-1-16S** | 334565 | 266205 | 265596 | 265846 | 204210 | 199658 | 60% |
| **M1-R3-T2-U-2-16S** | 391503 | 308982 | 308026 | 308294 | 236878 | 231449 | 59% |
| **M1-R3-T2-P-C-16S** | 302462 | 240121 | 238983 | 239371 | 213719 | 211968 | 70% |
| **M1-R3-T2-P-F-16S** | 262400 | 23372 | 22643 | 22989 | 21584 | 21488 | 8% |
| **M1-R3-T2-W-C-16S** | 252945 | 205155 | 204083 | 204870 | 128835 | 128619 | 51% |
| **M1-R3-T2-W-F-16S** | 397763 | 312142 | 310410 | 311526 | 265359 | 264191 | 66% |
| **M1-R4-T1-C-1-16S** | 461871 | 391647 | 391038 | 391044 | 315181 | 294626 | 64% |
| **M1-R4-T1-C-2-16S** | 743702 | 634522 | 633195 | 633064 | 490760 | 441999 | 59% |
| **M1-R4-T1-U-1-16S** | 410114 | 347236 | 345941 | 345764 | 294687 | 288182 | 70% |
| **M1-R4-T1-U-2-16S** | 579894 | 480455 | 478718 | 478612 | 406843 | 401851 | 69% |
| **M1-R4-T1-P-C-16S** | 542846 | 435480 | 434246 | 434276 | 387267 | 383259 | 71% |
| **M1-R4-T1-P-F-16S** | 407146 | 315326 | 313197 | 314284 | 288554 | 284710 | 70% |
| **M1-R4-T1-W-C-16S** | 346834 | 286409 | 284936 | 285487 | 204431 | 203339 | 59% |
| **M1-R4-T1-W-F-16S** | 280841 | 234465 | 231528 | 233030 | 205443 | 204412 | 73% |
| **M1-R4-T2-C-1-16S** | 768437 | 674416 | 673522 | 673901 | 545930 | 508096 | 66% |
| **M1-R4-T2-U-1-16S** | 400751 | 326204 | 325249 | 325220 | 265469 | 261315 | 65% |
| **M1-R4-T2-U-2-16S** | 413440 | 339384 | 338014 | 338467 | 265937 | 261801 | 63% |
| **M1-R4-T2-P-C-16S** | 419950 | 352357 | 351609 | 351949 | 332789 | 331917 | 79% |
| **M1-R4-T2-P-F-16S** | 1558165 | 1267723 | 1262107 | 1266263 | 1131827 | 1121758 | 72% |
| **M1-R4-T2-W-C-16S** | 489752 | 395719 | 394385 | 394903 | 338436 | 336616 | 69% |
| **M1-R4-T2-W-F-16S** | 353823 | 288667 | 286427 | 287669 | 248425 | 246136 | 70% |
| **M1-BLANK-16S** | 152760 | 130281 | 129501 | 129916 | 78068 | 77181 | 51% |

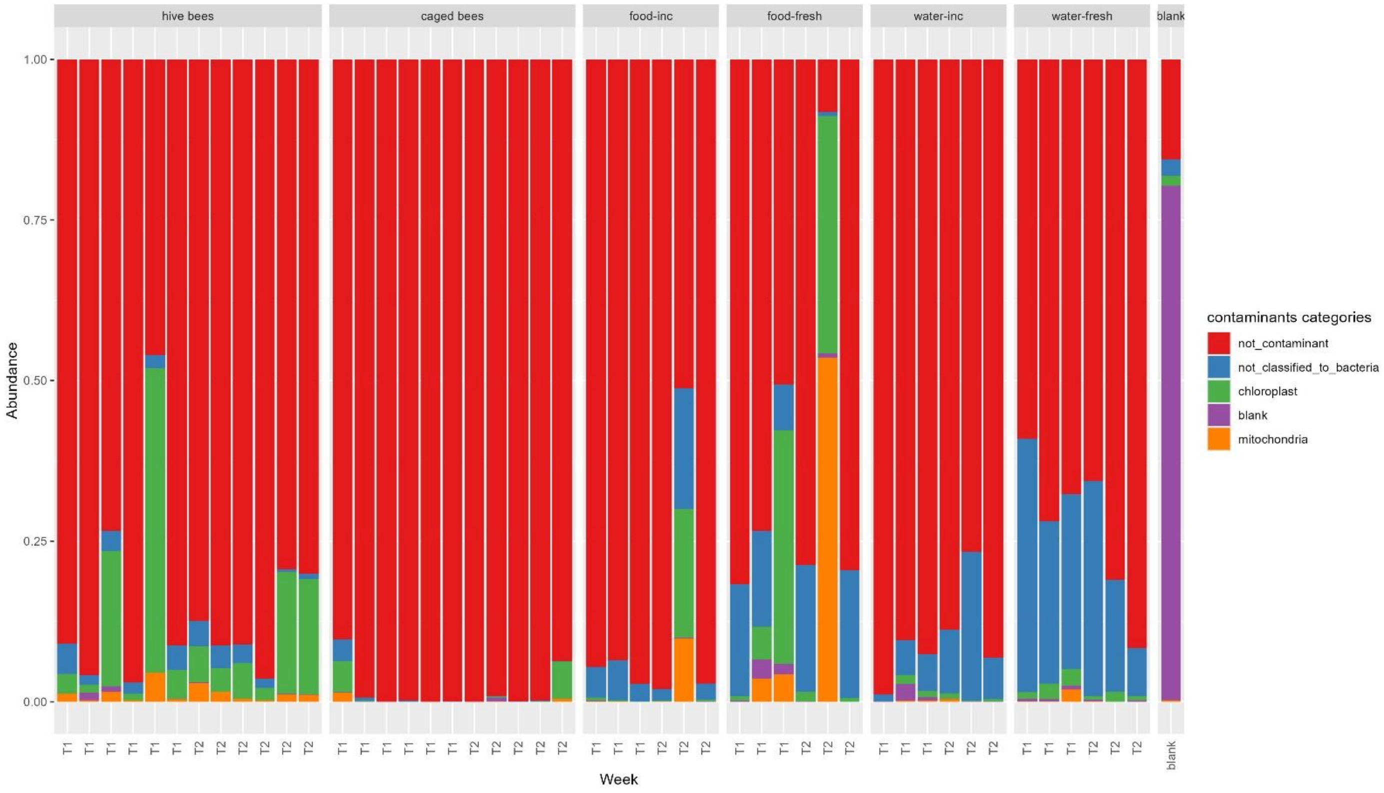


Fig. S1. Contaminant sources in the samples. 'inc' – food and water samples after 24 h in the incubator, 'fresh' – freshly prepared

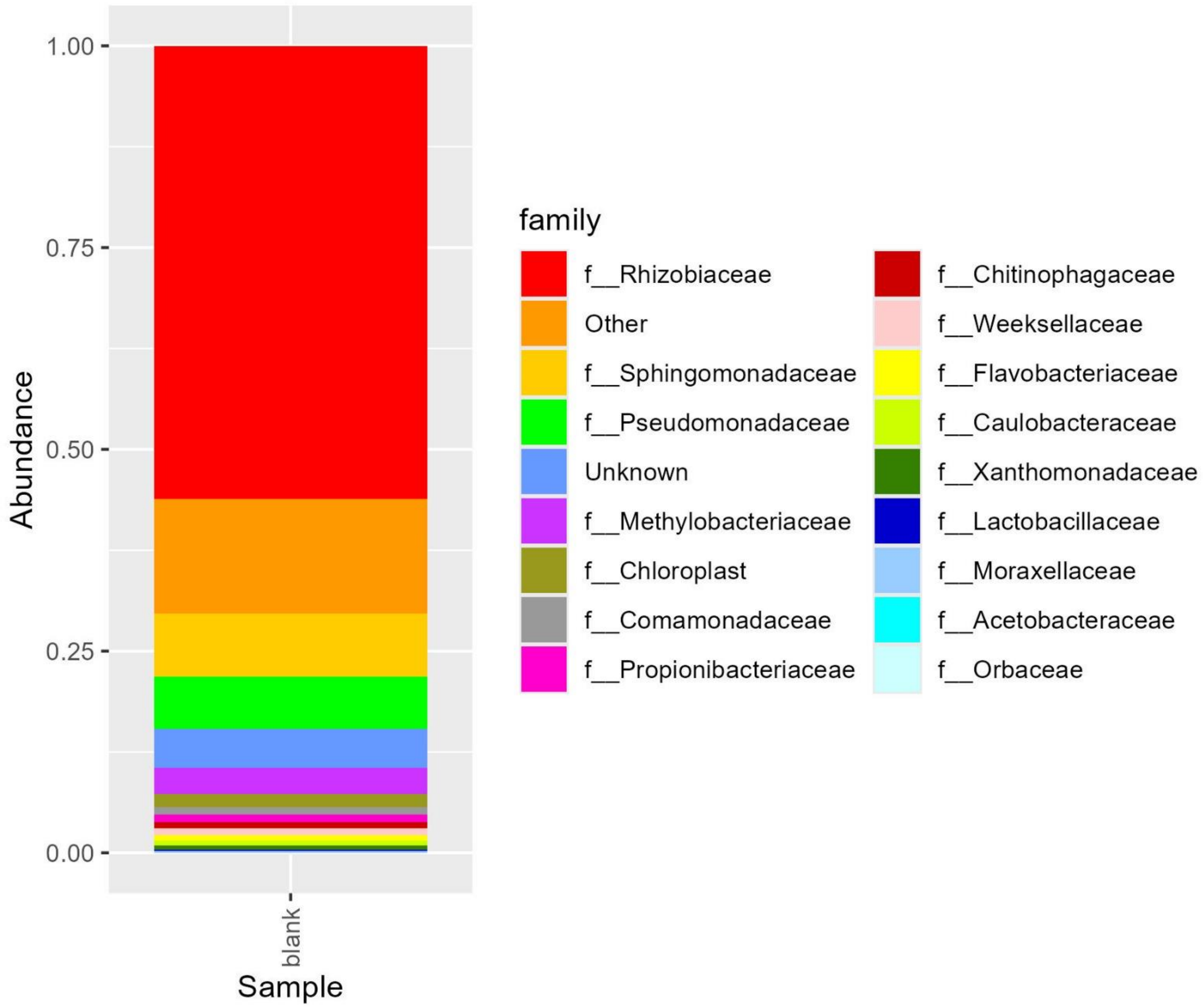


Fig. S2 Composition of the blank sample aggregated to family level.

Table S3 Divergence test results. Model: linear model, fixed factors: source, timepoint and their interaction

| | Df | Sum of squares | Mean square | F value | Pr(>F) |
|---|---|---|---|---|---|
| **source** | 1 | 2.88013 | 2.88013 | 49.6282 | 1.05e-06 |
| **timepoint** | 1 | 0.02042 | 0.02042 | 0.3519 | 0.5601 |
| **source:timepoint** | 1 | 0.08970 | 0.08970 | 1.5457 | 0.2289 |
| **residuals** | 19 | 1.10265 | 0.05803 | | |

Table S4. CCA model summary for bees samples

a. Model summary

| | Inertia | Proportion | Rank |
|---|---|---|---|
| Total | 3.2682 | 1.0000 | |
| Constrained | 0.7354 | 0.2250 | 3 |
| Unconstrained | 2.5327 | 0.7750 | 19 |

b. Eigenvalues for constrained axes

| CCA1 | CCA2 | CCA3 |
|---|---|---|
| 0.3995 | 0.2314 | 0.1046 |

c. Eigenvalues for unconstrained axes (8 of 19 unconstrained eigenvalues are shown):

| CA1 | CA2 | CA3 | CA4 | CA5 | CA6 | CA7 | CA8 |
|---|---|---|---|---|---|---|---|
| 0.5427 | 0.41793 | 0.3801 | 0.2081 | 0.1490 | 0.1321 | 0.1104 | 0.0917 |

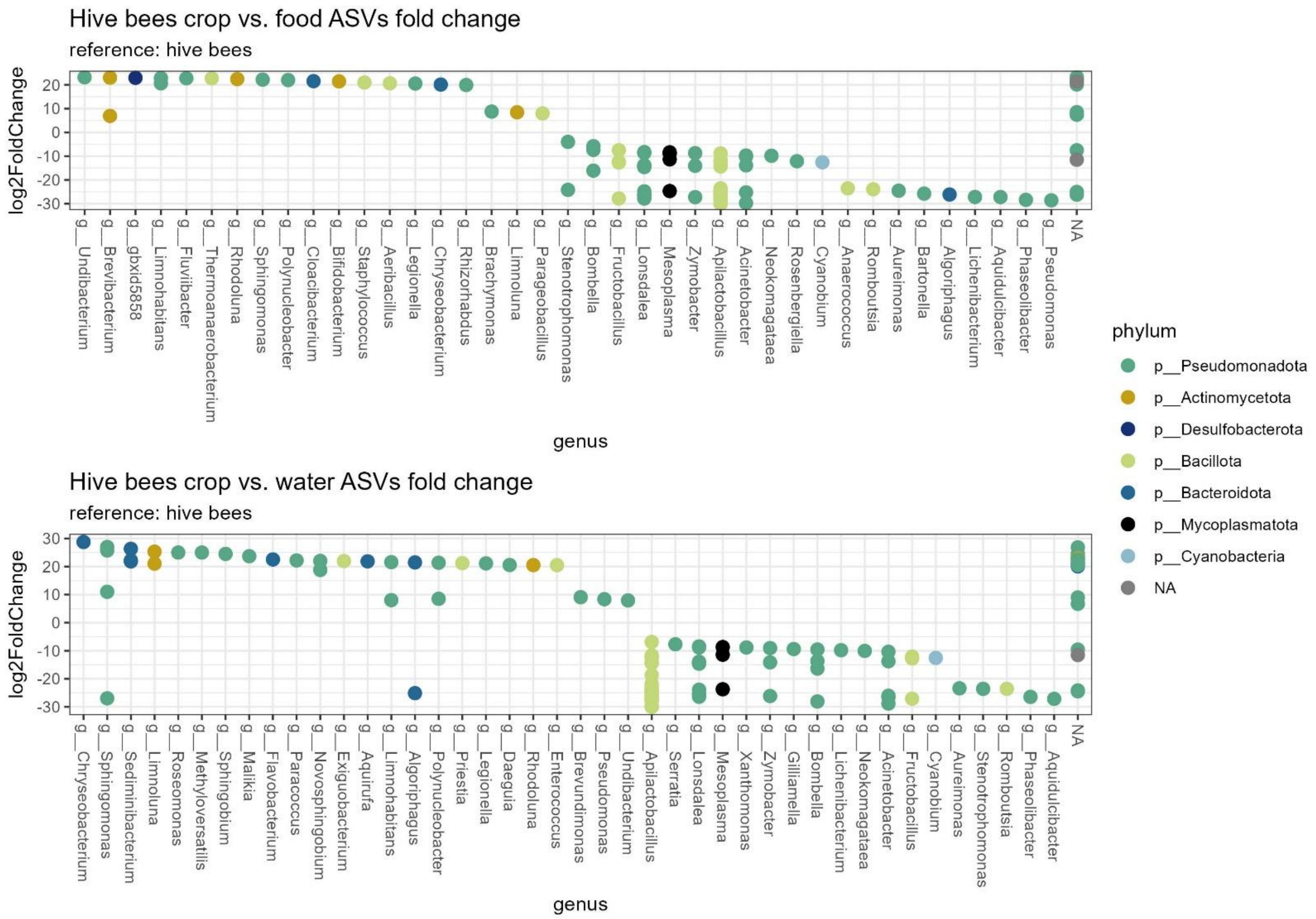


Fig. S3. ASVs fold change (analyzed with DESeq2) to identify bacteria characteristic for food (top panel) or water (bottom panel) samples rather than the natural bee crop microbiota (hive bee crop samples). Only the variants with a significant ($p<0.05$) fold change between the groups are plotted.
